# Comparison of potential plant-pollinator network structure for 3 seed mixes (lawn, roadside, and pollinator-planting) in western NY, USA

**DOI:** 10.1101/2020.10.29.352906

**Authors:** Briana Burt, Kristina Chomiak, Ibrahim Cisse, Aaron Paratore, Kaitlin Stack Whitney

**Affiliations:** Environmental Sciences Program, College of Science, Rochester Institute of Technology 84 Lomb Memorial Drive, Rochester NY 14623

## Abstract

There is growing concern, locally and globally, about the health of pollinating insects and their decreasing abundance and diversity. While roads may also be contributing to insect pollinator declines (roads can contribute to habitat fragmentation and habitat destruction), roadsides may provide opportunities for pollinating insect conservation. Yet to use these areas to support local pollinating insects, we need to understand which plants will support wild pollinators, especially of conservation concern. To that end, we researched the potential plant-pollinator networks of three existing seed mixes in western New York (USA) – a roadside seed mix, a pollinator-friendly planting mix, and a lawn seed mix. We used publicly available information and built bipartite graphs to show the resulting networks. The pollinator-friendly seed mix supported the most pollinating insects overall and taxa of conservation concern. Yet the roadside mix, with the same species richness as the lawn seed mix, supported a different network based on the plants in the mix. Our results inform which particular plant species in existing seed mixes in western New York can support wild pollinating insect species of concern in the region. Additionally, our results show potentially how roadside and lawn plantings may be altered to support a broader network of pollinating insects.

## Introduction

There is growing concern, locally and globally, about the health of pollinating insects and their decreasing abundance and diversity. Among other stressors, the loss of habitat from development has contributed to the widespread decline of pollinators throughout the nation (NRC 2007). Urban development and habitat fragmentation have also contributed to the decline of pollination services (Cunningham 2000). Habitat fragmentation has additional impacts on biodiversity, seed dispersal, and nutrient cycling from edge effects (Cuke and Srivastava 2016; Malmivaara-Lamsa, et al. 2008). The effects of habitat fragmentation apply to both insect and avian pollinators through habitat loss and reduction in available food sources.

Roads may also be contributing to insect pollinator declines. Roads can contribute to habitat fragmentation and habitat destruction (Hunter 2002). Studies on road impacts have increased in part due to their ubiquity. Roads have grown to encompass approximately 1% of the land area in most developed countries. Additionally, they are estimated to have direct ecological effects on 20% of the land area in the United States (Forman 2000). Though less than 1% of U.S. land is paved roads, an estimated 15–20% of total land area in the US is directly affected ecologically by roads through habitat fragmentation and road-avoidance by wildlife from the noise and speed (Dargas et al. 2016; as reviewed by Forman &Alexander 1998). For pollinating insects as small animals, habitat fragmentation from roads can be ruinous by altering key aspects of their habitat, in addition to immense loss of habitat (as reviewed by Coffin 2007). In the United States, wild pollinators are declining due to habitat destruction, invasive species, and disease by way of road construction (Kluser et al. 2007).

While roads may be detrimental to conservation efforts, the areas along them may provide a unique opportunity for wildlife conservation by harnessing them as habitat. Roadside conservation interventions may be a way to increase native plant and insect populations such as monarch butterflies (Pleasants 2017, Stack Whitney 2017). Roadsides have the potential to be good habitat for pollinators when well managed. For example, restored roadsides support an abundance of bees and have higher species richness compared to weedy roadsides (Hopwood 2008).

Roadsides are just one of several anthropogenic habitats, such as lawns, that may be used for pollinator insect habitat. However, the efficacy of these habitats and the plants in them is still under research. There are some promising results to date. Native wild pollinating insects, especially bees, have preferred host plants, and specialists cannot switch to use non-preferable plants in the absence of those plants (Angelella et al. 2019). Yet some wild pollinating insects are generalists and may be able to use a variety of plants, including those found in anthropogenic habitats. There is a need to understand which specific plants can potentially support wild pollinators in our area. This information can also be used to inform whether additional plant species could be added to the existing seed mixes to support additional or more diverse pollinating insects in the region.

Our objective was thus to examine the potential plant-pollinating insect interaction networks of three local seed mixes in Western NY – a lawn seed mix, a roadside seed mix, and a pollinator-friendly planting seed mix. We expected that the pollinator-specific seed mix would support the highest number of pollinating insects. Our goal was to determine which local pollinating insects use the plant species in each seed mix, then determine the conservation status of each and to understand which if any mixes support species of conservation concern in the region. The results will inform how roadsides compare to other plantings as potential habitat to contribute to pollinating insect conservation and potentially serve to provide recommendations on how to improve roadside seed mixes for land managers who want to use roadsides as habitat.

## Methods

### Seed mix selection

We chose three local seed mixes that are used in western upstate New York. We included one from each habitat type (roadside, lawn, pollinator-friendly planting) to understand the potential plant-pollinating insect interaction network of different plantings in the region. The first was a roadside seed mix for New York (Appendix 1A, New York Department of Transportation 2014). The second was a lawn seed mix for New York (Appendix 1B, New York Department of Transportation 2014). The third seed mix was a pollinator-friendly planting seed mix sold commercially to homeowners in the greater Rochester area (Appendix 1C, Will 2018). We noted the date of each seed mix, as the mixtures may change in the future.

### Plant conservation information

Using the plant species list for each seed mix, we then researched the origin of each plant species (whether native to North America and New York) and known conservation status (both in New York and globally). This was compiled from publicly available sources on the internet. We used the New York Flora Atlas (Weldy et al. 2019), as well as information from the United States Department of Agriculture Natural Resource Conservation Service PLANTS Database (USDA, NCRS 2019), Missouri Botanical Garden Plant Finder (Missouri Botanical Garden Plant Finder 2019) and Xerces Society for Invertebrate Conservation (Adamson et al. 2017). We then made tables with the NY rank, global rank, endangered or threatened status, and native or naturalized status for each plant species and seed mix (Table 1, Table 2, and Table 3).

**Table 1.**
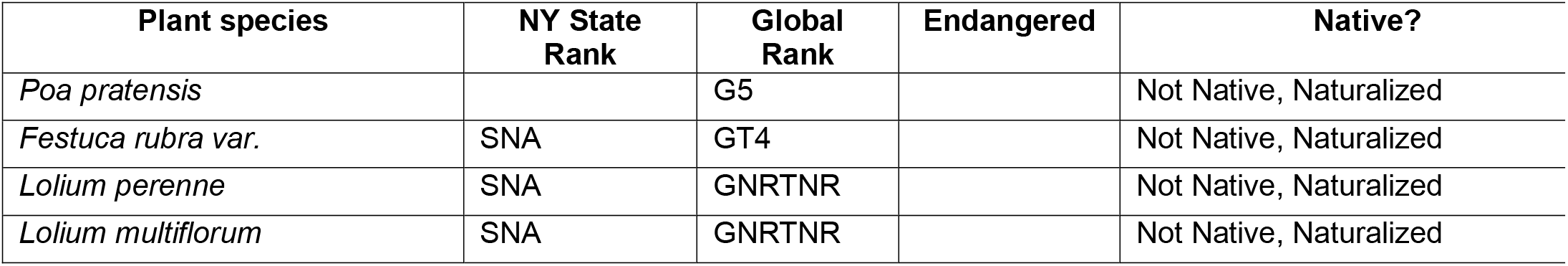
Conservation status of the plants in the lawn seed mix at state and global scales.

**Table 2.** Conservation status of the plants in the general roadside seed mix at state and global scales.

| Plant species | NY State Rank | Global Rank | Endangered | Native? |
| --- | --- | --- | --- | --- |
| *Festuca rubra* var. | SNA | GT4 |  | Not Native, Naturalized |
| *Lolium perenne* | SNA | GNRTNR |  | Not Native, Naturalized |
| *Lolium multiflorum* | SNA | GNRTNR |  | Not Native, Naturalized |
| *Trifolium repens* | SNA | GNR |  | Not Native, Naturalized |

**Table 3.**
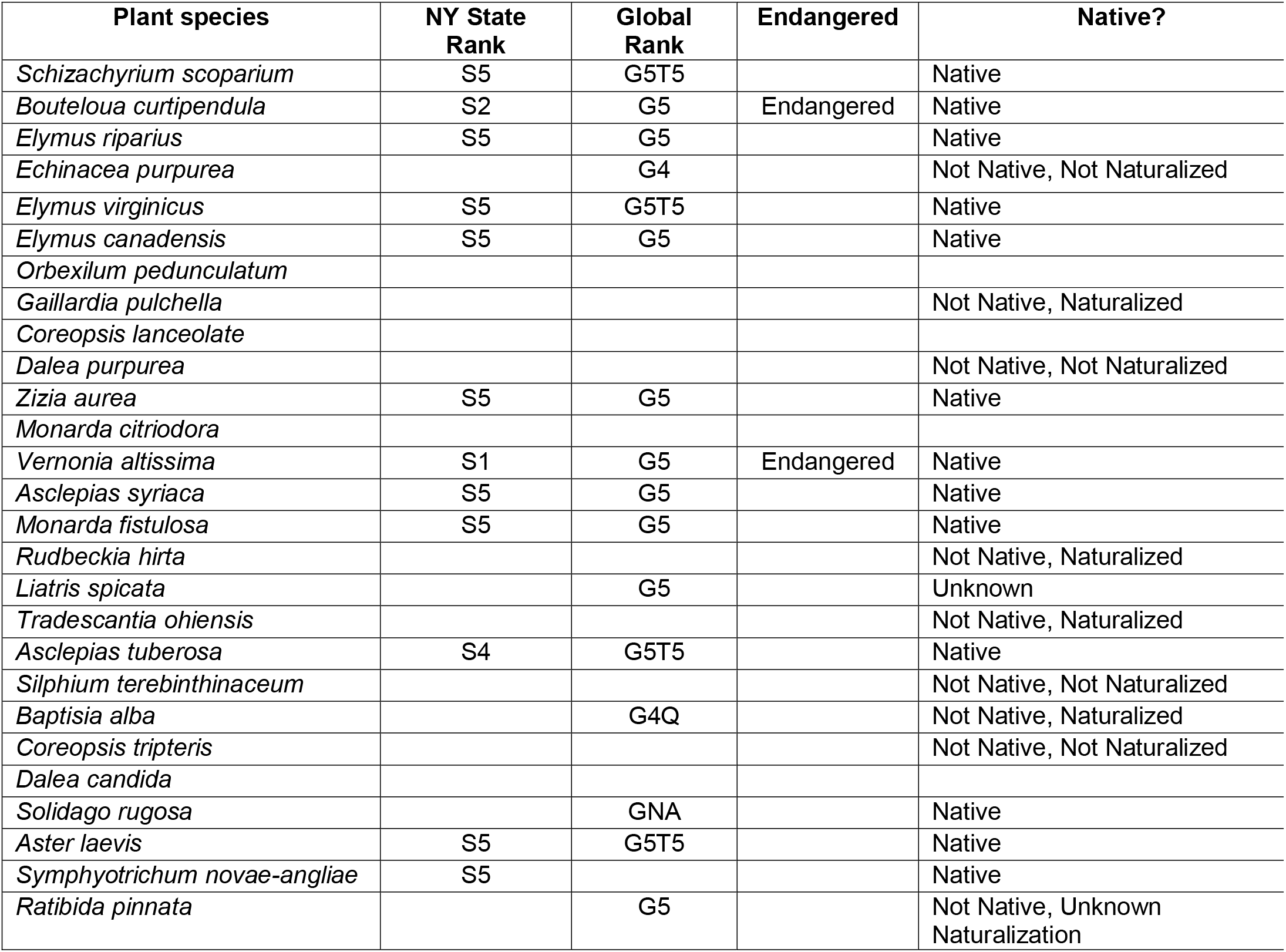
Conservation status of the plants in the Seneca Park Zoo Society pollinator friendly seed mix at state and global scales.

### Pollinating insect visitor research

To investigate which pollinating insects in western New York may visit the plant species in each of the three mixes, we used publicly available information about which organisms are known to pollinate or otherwise visit the plants. We triangulated information about each plant species from several sources to ensure the broadest possible description of known visitors in the region. Our sources included the Illinois Wildflower website (Hilty 2017), Xerces Society for Invertebrate Conservation (Adamson et al. 2017; Sheperd et al. 2008), Lady Bird Johnson Wildflower Center (NPIN 2013), and the New England Wild Flower Society (Haines et al. 2011). We also searched each plant species in the Web of Science journal article database using the search terms “[*scientific name of plant]*” AND pollinat* (Clarivate 2019). Additionally, we also checked for any other information using the Google search database with the search terms “[*name of plant]*” AND “pollinator” OR “insect association” OR “faunal association” (Google Search 2019).

### Pollinating insect visitor attributes

Once we had a list of all the known pollinating insect visitors for each plant species in the three mixes, we looked up conservation status information about each insect taxonomic group. We used publicly available information online in BugGuide (BugGuide 2018), Butterflies and Moths of North America (Lotts and Naberhaus 2017), the National Wildlife Federation database sourced from United States Fish and Wildlife Service (NWF 2019), and the Great Pollinator Project (Matteson 2019). This was in part to confirm that they were found in western NY; any pollinating insects found in the previous that were not found in western NY were removed. All bee information was obtained from the Cornell Bees of New York “species list – bees of New York” (Pollinator Network @ Cornell 2019). We also used these sources to investigate the conservation status of each pollinating insect taxa, as well as information from the New York State Department of Conservation list of endangered, threatened, and special concern fish and wildlife species (NYSDEC 2015).

### Plant-pollinator network mapping

To visualize the network connections of the pollinating insects and the plant species, we created bipartite graphs in R statistical software (R Core Team 2017). In these plots, a line connects each plant species to each pollinating insect species known to visit (Figure 1A, 1B, and 1C). To do this, we first had to transform the plant and insect information into a spreadsheet with binary inputs. We scored each relationship (a plant species supporting an insect species) as a 1. All other insect species were scored a 0. The Igraph package was used and all instructions were taken from the documentation associated with the package (Csardi and Nepusz 2006).

**Figure 1:**
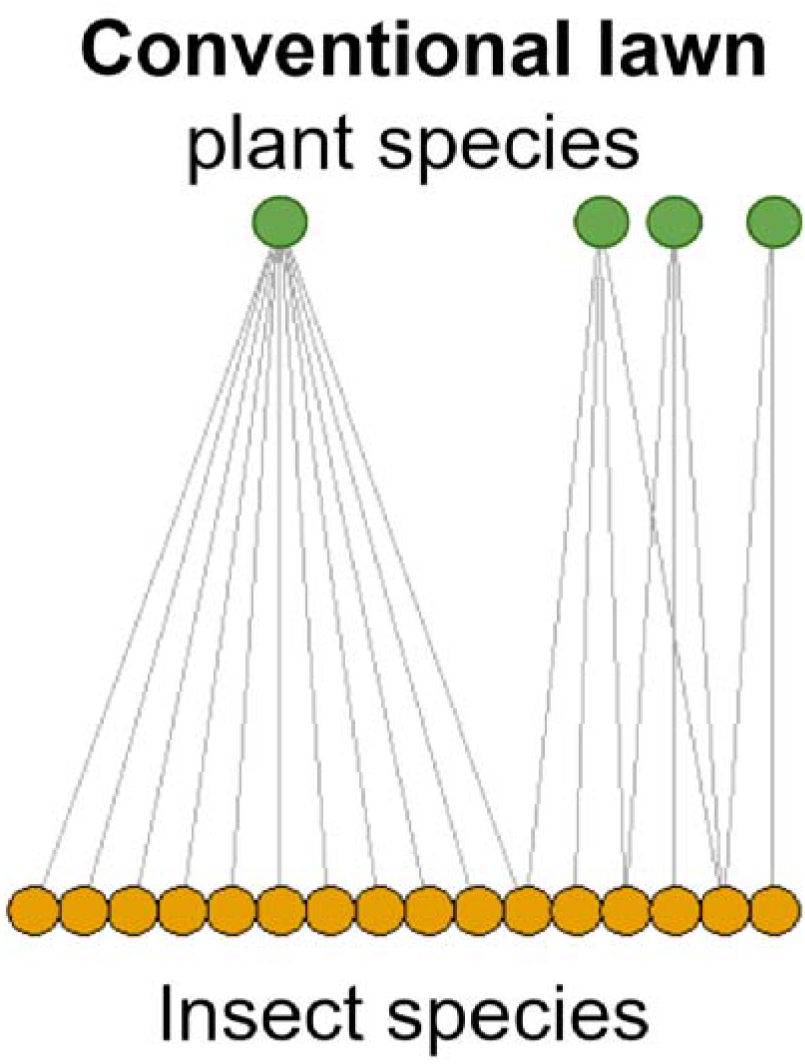
Bipartite graph of the plant-pollinator network for the lawn seed mix. There are 4 plant species that could potentially support 16 species of pollinating insects in western NY.

## Results

### Plant attributes

We found that the lawn seed mix contained 4 species, all of which are not native but are naturalized to the area and the country (Table 1, Appendix 1A). The same is true for the general roadside seed mix; it has four species in the seed mix. The only difference being that the lawn mix has *Lolium* spp. (Italian ryegrass) but in the roadside seed mix it is replaced with *Trifolium repens* (white clover) (Table 2, Appendix 1B).

The commercially available pollinator-friendly seed mix has many more plant species, this particular mix has 27 species (Appendix 1C). Compared to the lawn and roadside seed mixes, many of the plants are native (Table 3). Additionally, several of the plant species are considered to be rarer and of conservation concern.

The mixes also varied in the proportion of plants in the mix that are flowering and may provide nectar resources. The roadside seed mix is composed of three grasses and one forb (Table 2). The lawn seed mix is composed of four forbs (Table 1). Conversely, the pollinator-friendly seed mix is made up of a majority of flowering and nectar-producing plant species (Table 3).

### Plant-pollinator networks

We found stark variation in the structure and composition of the plant-pollinator networks from the three plant seed mixes we examined and their pollinating insect visitors known in western New York. Due to the size, please note that the complete tables are all in a workbook in the supplemental information (Supplemental Table 1).

We found that the four grass species in the lawn seed mix could potentially support 16 species of pollinating insects in the region (Figure 1, Supplemental Table 1). The majority of the insect pollinators supported were Lepidoptera, including skippers, moths, and butterflies (Supplemental Table 1). *Poa pratensis* supported the most insects of the species in the mix.

The roadside seed mix, which included three grass species and one flowering forb, could potentially support 77 species of known pollinating insects in western NY (Figure 2, Supplemental Table 1). The majority of the increase in pollinating insects potentially supported comes from the addition of white clover (*Trifolum repens*) to this seed mix, which has the ability to support many different insect visitors. White clover supports butterflies, flies, and long-tongued wild bees. Butterflies and moths are also potentially supported by this seed mix (Supplemental Table 1).

**Figure 2:**
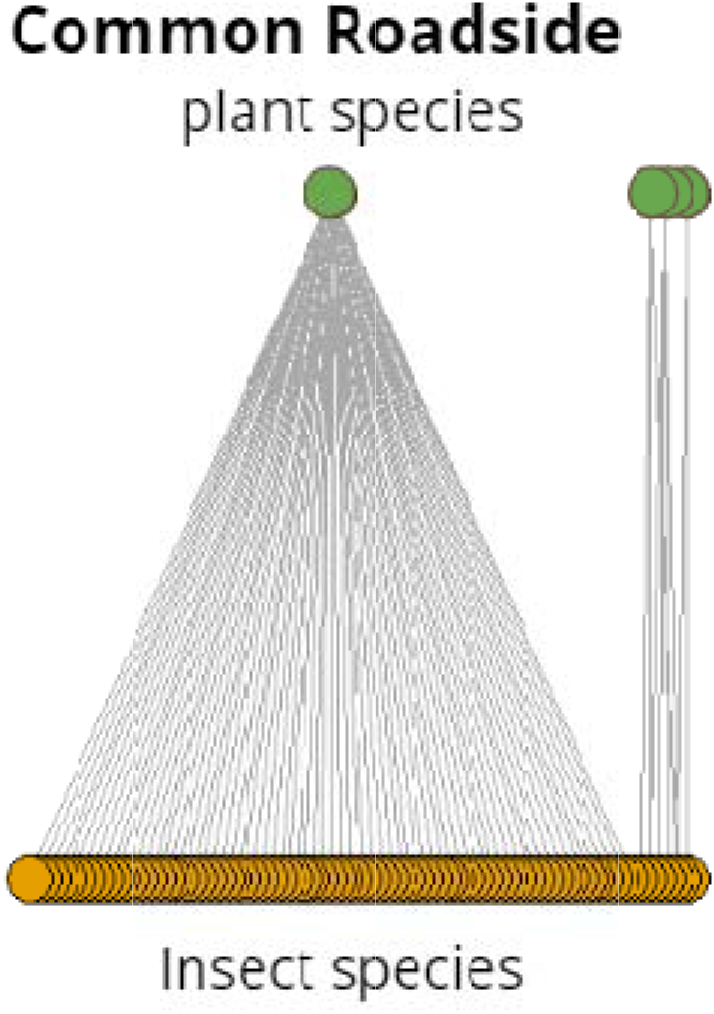
Bipartite graph of the plant-pollinator network for the General Roadside seed mix. There are 4 plant species that could potentially support 77 species of pollinating insects in western NY.

The structure of the potential plant-pollinator network from the pollinator-friendly seed mix available for residential areas is much richer (Figure 3). Both the number of insect species found in western New York that may be supported by the mix and the connection nodes between the plant and insect species is much higher than that from the lawn or roadside seed mix. With 27 plant species, this seed mix could potentially support 520 species of pollinating insects whose known range includes western NY (Figure 3, Supplemental Table 1). Importantly, this includes potentially supporting 9 insect species listed as endangered in New York and 53 species which are known to be declining in New York (NYSDEC 2015).

**Figure 3:**
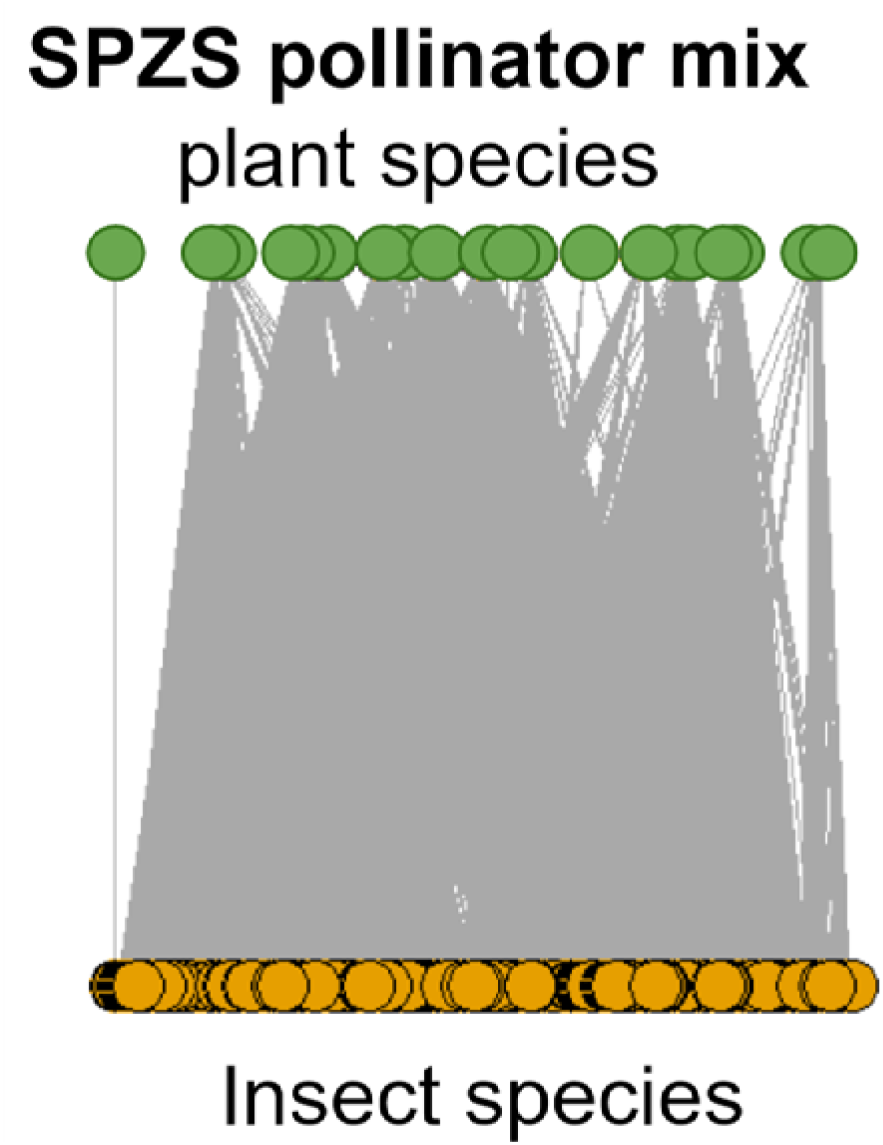
Bipartite graph of the plant-pollinator network for the pollinator friendly seed mix available from the Seneca Park Zoo Society. There are 27 plant species that could potentially support 520 species of pollinating insects in western NY.

## Discussion

Overall, we found that existing seed mixes for lawns and roadsides do not contain many plant species, compared to a pollinator-friendly seed mix. In turn, the pollinator planting seed mix supported a significantly higher number of pollinating insect species than either the lawn or roadside seed mix. This is due to the number of flowering nectar plants that supported many pollinating insects, both an increased number of species and diversity of types of insects. This result is in line with previous research which has shown that there is a positive correlation between pollinator abundance and wildflower bloom counts (e.g. Angelella et al, 2019).

However, our results also demonstrate that even seed mixes with relatively few plant species in them can support an increased number and diversity of insect visitors with the addition of flowering plants that provide nectar and pollen. The difference in the pollinating network supported by the roadside seed mix by the inclusion of one forb (*T. repens*, white clover), demonstrates how the specific plant species and species diversity, not just species richness, in the seed mix significantly impacted the network that may potentially be supported in western New York.

Our results have critical implications for conservationists and planners in the region. When designing seed mixes and planting projects intended to enhance wild pollinators and wild pollination, it must be done with an understanding of which insect species may be supported. Our results demonstrate that existing lawn and roadside seed mixes are limited in their ability to support wild insect pollinators.

Our findings are also important for landscape ecologists and other ecology researchers studying roadside and lawn habitats. Many of the services and species found in previous research in these habitats are likely due to volunteer and weed species, not the species in the plant mix itself. This makes management choices critical to understand – and explicit discussion of the plant biodiversity in the studied habitats.

It is important to point out that this is a representation of only pollinating insects that are being supported by the plants and not all organisms, therefore it is an underestimate of the actual number of organisms supported by any of the seed mixes. The plant-pollinator networks presented in this research represent the total possible networks found with the resources currently available. One limitation to this study included the fact that many insects are not well studied or recorded. This made specific insects difficult to research or may have caused omission of potentially important insects. With thousands of potential pollinating insects in New York State, this list is biased towards the most well-studied. Unfortunately, this lack of information also limits the available knowledge of whether an insect species is declining.

Ultimately, as urbanization continues, the ecological implications for pollinating insects will be magnified as roadways and other impervious surfaces from development continue to expand. Overall, our research shows the potential benefit diverse, intentional pollinator-friendly plantings for wild pollinator conservation, especially relative to traditional lawns and current roadside plantings. Yet this does not mean that lawn and roadside seed mixes must remain as they are now. Our findings can be used to determine which plants could potentially be added to future roadsides or lawns to benefit wild pollinating insects of conservation concern in our region.

## Appendix 1 Seed mix lists

## Appendix 1A: Roadside seed mix

The plant species in the “general roadside seed mix” in the New York State Department of Transportation Standard Specifications report (NYSDOT 2014).

| **Latin Name** | **Common Name** |
| --- | --- |
| Festuca rubra | Fine fescue |
| Lolium perenne | Perennial ryegrass |
| Lolium multiflorum | Annual ryegrass |
| Trifolium repens | White clover |

## Appendix 1B: Lawn seed mix

The plant species in the “lawn seed mix” in the New York State Department of Transportation Standard Specifications report (NYSDOT 2014).

| **Latin Name** | **Common Name** |
| --- | --- |
| Poa pratensis | Kentucky Bluegrass |
| Festuca rubravar. | Fine Fescue |
| Lolium perenne | Perennial Ryegrass |
| Lolium multiflorum | Annual Ryegrass |

## Appendix 1C. Pollinator-planting seed mix

The plant species in the pollinator-friendly planting residential mix from the Seneca Park Zoo Society (Will, 2018).

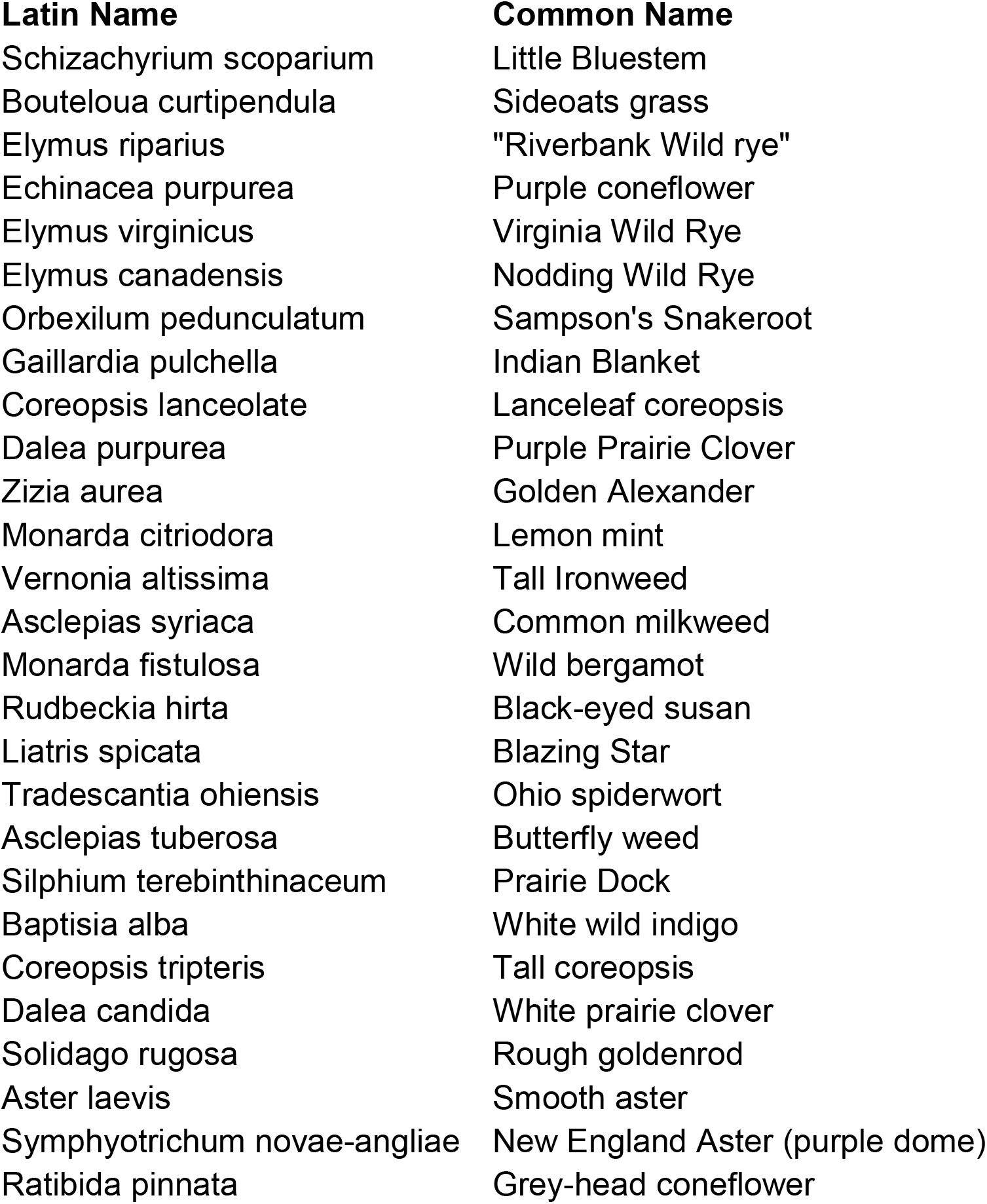

